# Evolution-informed forecasting of seasonal influenza A (H3N2)

**DOI:** 10.1101/198168

**Authors:** Xiangjun Du, Aaron A. King, Robert J. Woods, Mercedes Pascual

**Affiliations:** Department of Ecology and Evolution, University of Chicago, Chicago, IL, USA; Departments of Ecology & Evolutionary Biology and Mathematics, University of Michigan, Ann Arbor, MI, USA; University of Michigan Health System, University of Michigan, Ann Arbor, MI, USA; The Santa Fe Institute, Santa Fe, NM, USA

## Abstract

Inter-pandemic or seasonal influenza exacts an enormous annual burden both in terms of human health and economic impact. Incidence prediction ahead of season remains a challenge largely because of the virus’ antigenic evolution. We propose here a forecasting approach that incorporates evolutionary change into a mechanistic epidemiological model. The proposed models are simple enough that their parameters can be estimated from retrospective surveillance data. These models link amino-acid sequences of hemagglutinin epitopes with a transmission model for seasonal H3N2 influenza, also informed by H1N1 levels. With a monthly time series of H3N2 incidence in the United States over 10 years, we demonstrate the feasibility of prediction ahead of season and an accurate real-time forecast for the 2016/2017 influenza season.

**SUMMARY:** Skillful forecasting of seasonal (H3N2) influenza incidence ahead of the season is shown to be possible by means of a transmission model that explicitly tracks evolutionary change in the virus, integrating information from both epidemiological surveillance and readily available genetic sequences.

## INTRODUCTION

Inter-pandemic or seasonal influenza exacts an enormous public health burden around the globe, with an average of about 1 billion cases, including 3 to 5 million cases of severe illness and 250 000 to 500 000 deaths annually (*1*). Since its first occurrence in 1968, seasonal H3N2 influenza has continually circulated in the human population, and is currently the major cause of seasonal influenza morbidity and mortality (*2*). The sustained ‘success’ of influenza viruses responsible for seasonal outbreaks stems from their ability to evolve and escape the immune system by modifying their surface proteins (*3*). Phylogenetic trees depicting evolutionary changes in (H3N2) influenza viruses illustrate rapid drift with successive and punctuated replacement of one antigenic type by another (*4*, *5*). The last decade has seen significant conceptual advances in the understanding of these phylogenetic patterns, enabled by computational and statistical advances at the interface of transmission dynamics and virus evolution (*5*-*12*). There is now considerable interest in translating these conceptual advances into actual prediction at the population level that would inform the update of vaccines and epidemic preparedness.

The challenge of influenza prediction has progressed largely along two separate tracks. On the one hand, there are computational methods based on phylogenies and mutation patterns in the surface protein hemagglutinin (HA), whose goal is to predict evolutionary change (*13*-*17*). The resulting predictions of successful lineages and their relative frequencies for the future season do not, however, provide precise information on absolute incidence. On the other hand, mathematical models describing the transmission dynamics of influenza viruses allow real-time incidence forecasts of influenza-like illness (ILI) (*18*, *19*). With data assimilation methods (*20*), these models must be fitted within each season because of season-to-season viral evolution (*21*-*25*). In other words, with such models, the fact that one needs to wait until the outbreak starts limits the lead time of epidemiological prediction.

In this study, we bridge the gap between these two approaches and propose an epidemiological model specifically for seasonal H3N2 influenza that incorporates information on the evolutionary change of the virus. The resulting model is sufficiently parsimonious that parameter estimation based on retrospective surveillance records is possible. A novel feature is its use of an evolutionary index of virus innovation constructed using readily available sequence data. The goal is to generate H3N2 incidence forecasts before the season begins, significantly earlier than what is currently possible. We illustrate two model formulations for H3N2 in the United States (US) between 2002 and 2016, and produce a forecast for the upcoming 2016/2017 influenza season. We emphasize prediction of interannual disease risk rather than finer-scale outbreak timing during the season; in other words, we seek to forecast whether or not the upcoming season will be anomalously large or small. Timing itself has been the target of existing within-season prediction efforts, which are better suited for this purpose and could be applied in tandem with our approach.

## RESULTS

The monthly incidence variability from 2002 to 2016 for the whole US is shown in Figure 1A for reported cases of influenza type A, for subtypes H3N2 and H1N1 (including both seasonal H1N1 and pandemic H1N1), as well as type B. The time series are computed as the product of the ILI positive rate, the influenza positive rate, the subtype proportion, and the US population size. Thus, incidence data are extrapolated to the US population from outpatients in a network of healthcare providers, with un-typed influenza specimens assigned to H3N2, H1N1 and B respectively based on the proportions from the US surveillance system (see Materials and Methods for details). The temporal variability of H3N2 exhibits seasonal outbreaks whose size varies considerably from one year to the next. This interannual variability can result from epidemiological processes such as the loss of immunity to a specific variant of the virus (*26*), but also to a large degree, from the antigenic evolution of the virus (*27*) and the combined and complex interactions of the two (*28*, *29*).

**Figure 1.**
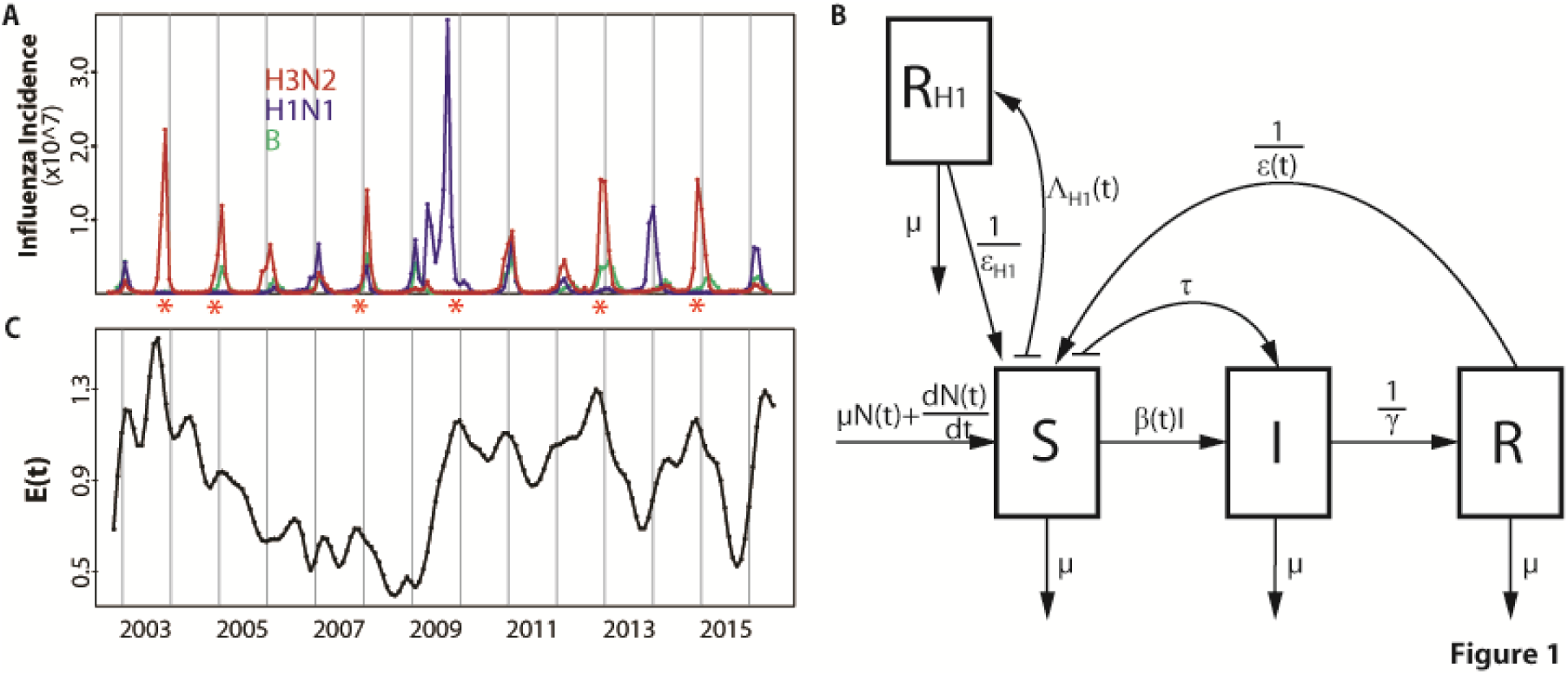
Data and model. **(A)** Monthly influenza incidence data for the US between October 2002 and June 2016. Red, blue and green curves are for subtype H3N2, subtype H1N1 and type B respectively. Seasons with an antigenic cluster transition were marked with asterisks. **(B)** Diagram for the epidemiological model. A classical susceptible – infected – recovered-susceptible (SIRS) epidemic model was used to represent the population dynamics of H3N2 incidence. The SIRS model is a compartmental formulation that follows the number of individuals into three classes, for susceptible (*S*), infected (*I*) and recovered (*R*) individuals respectively. People die at a constant rate μ. *N*(*t*) is the population size and the birth of individuals was specified as 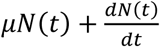 to capture the observed increase of the population. Susceptible individuals in *S* move to the *I* class after contact with an infective and transmission of the disease at rate *β* (*t*). This transmission rate includes a seasonal component, a dependency on the antigenic change of the virus, and environmental noise. Infected individuals eventually recover with an average infectious period of *γ* and move to the *R* class where they are protected by acquired immunity. Specific immunity is temporary and will be lost after an average latent period *ε* (*t*), with individuals in *R* returning to the *S* class. Parameter τ is the rate of external importation of H3N2 cases. An additional *R* _*H*1_ class was designed to track the protected population due to infection by H1N1. The rate of transition to the *R* _*H*1_ class is given by Λ _*H*1_(*t*), which depends on the observed incidence of H1N1 scaled to take into account the estimated reporting error. Individuals in the *R* _*H*1_ return back to the R class after an average latent period of ε _*H*1_. **(C)** Monthly evolutionary change *E*(*t*). The transmission rate *β* (*t*) in our first model incorporating evolutionary change depends on this evolutionary index. *E*(*t*) was calculated based on epitope sites of HA, as a weighted sum of normalized amino acid distances (hamming distances) between strains in month *t* and previous strains. Those distances were weighted by a decaying function back in time whose time scale was estimated as part of the model fitting. Details are described in the Evolutionary Index section of the Materials and Methods.

Before examining predictions of the ‘full’ model that considers both epidemiology and evolution, we evaluate the ability of different models, encapsulating different degrees of complexity, to retrospectively explain the temporal patterns in the data from 2002 to 2016. To establish a baseline against which to evaluate the full model, we begin with a simpler formulation for the population dynamics of H3N2 that describes influenza epidemiology and the seasonality of transmission but does not yet include evolutionary change (Fig. 1B). This basic model follows the structure of the well-known compartmental susceptible-infected-recovered-susceptible (SIRS) formulation which divides the population into classes for susceptible (non-immune), infected, and recovered (immune) individuals. For the purpose of model comparisons, we rely initially on the whole temporal extent of the data (2002-2016) to fit the models and infer parameter values. Figure 2A shows that simulations of the basic model reproduce the average seasonality of incidence but fail to capture its interannual variation.

**Figure 2.**
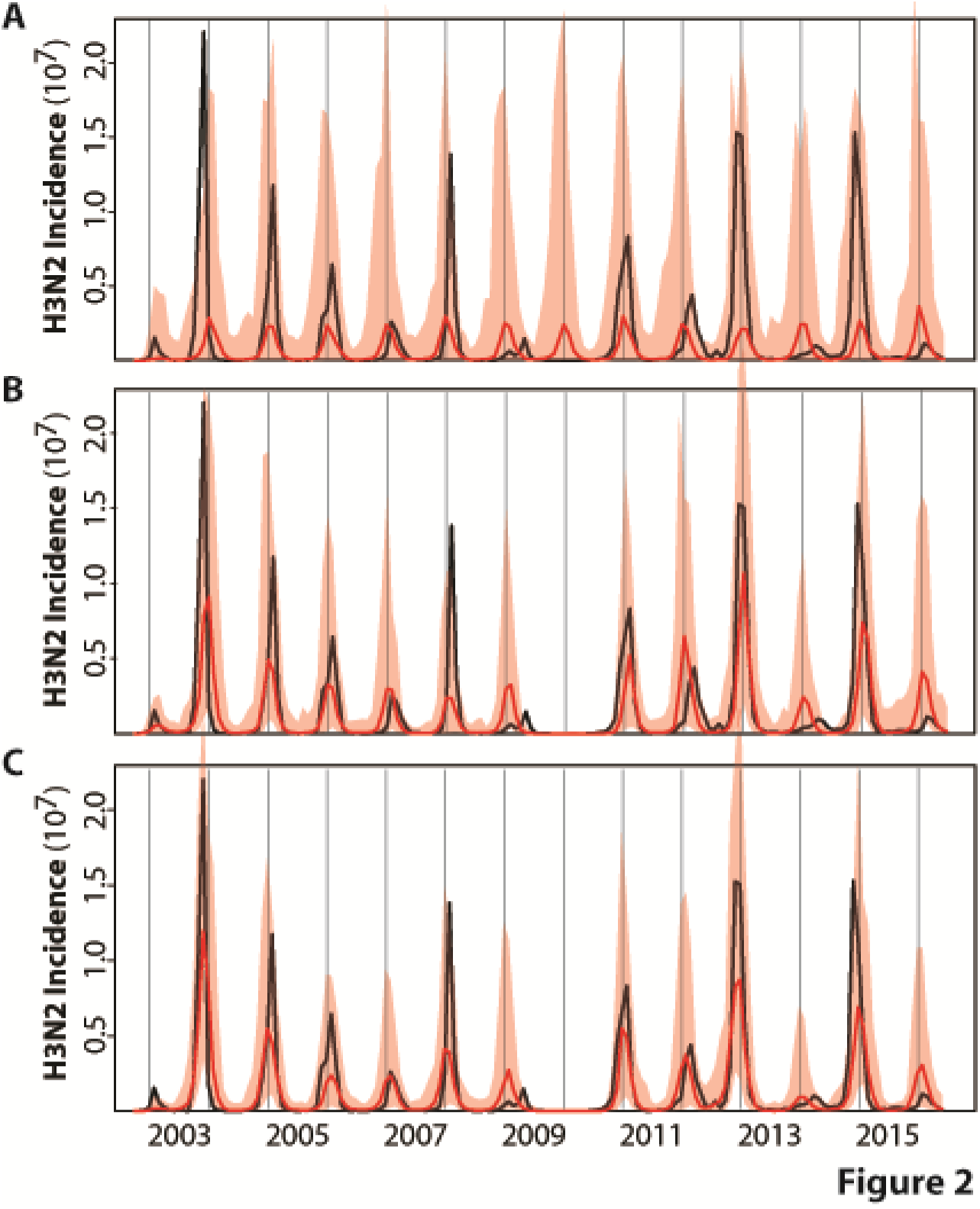
Illustration of the best model fits for the (**A)** basic, **(B)** continuous and **(C)** cluster models. See Table 1 for the specification and statistical comparison of the different models considered. Here, monthly simulations of the respective models with the MLE (Maximum Likelihood Estimates) parameters are shown for the median (in red) and 2.5-97.5% quantiles (shaded red) of 1000 simulations starting from estimated initial conditions in October 2002. For comparison, the observed monthly H3N2 incidence data for the US are shown in black. The basic model which incorporates only a fixed seasonality and no information on H1N1 in (A) fails to capture the temporal variability in the size of seasonal outbreaks; whereas the two models that include a dependence on the levels of H1N1 and on the evolutionary change of the virus (in a continuous fashion in B, and a discrete one in C) do represent this interannual variation.

Several variants of the epidemiological model were considered next, starting with the effect of temporal variation in the incidence of H1N1 (Fig. 1B). We find a significant negative correlation between annual incidence of H3N2 and H1N1 (*r* =-0.60, *p value* = 0.02), but no clear relationship between annual incidence of type B and H3N2 or H1N1 (*r* = 0.42 and *r* =-0.33, *p values* = 0.13 *and* 0.25, respectively). Given these findings and the observation that disease burden due to type B in humans is typically lower than that due to H3N2 or H1N1 in the US (*30*, *31*), we postulate an effect of H1N1, but not B, on the dynamics of H3N2, and do so as a covariate affecting the system as an external observed variable (in the ‘basic-H1’ model, Fig. S1A). We return to this simplification in the discussion. We also allow measurement error to differ between the summer (April 1^st^ to September 30 ^th^) and winter (October 1^st^ to March 31^st^) seasons, reasoning that the reporting rate is more variable outside the winter (transmission) season (*32*). (Hereafter, we refer to the year/year + 1 influenza ‘season’ as the period from July 1^st^ of the current calendar year to June 30 ^th^ of the following calendar year, for example, the 2008/2009 influenza season). Additionally, we relax the assumption of a linear dependence of the force of infection on the number of current infected individuals, allowing a nonlinear functional form and the potential for sub-exponential growth of the epidemic curve (in the ‘refined’ model, Fig. S1B). This functional form has been found effective in the modeling of a number of different infectious diseases, as a means of parameterizing processes operating at scales smaller than can be explicitly represented (*33*-*35*). Finally, we consider a version of the refined model in which the (non-parametric) periodic function in the transmission rate is replaced by a function of specific humidity (the ‘humidity’ model, Fig. S1C) (*21, 36, 37*). Although all these model variants improve the fit of the data (Table 1), they still fail to properly capture the interannual variability in incidence (Fig. S1). Of these purely epidemiological models, we retain the best, viz. the refined model, which includes H1N1, season-dependent measurement error, and sub-exponential epidemic growth, and turn next to whether the inclusion of an index of evolutionary change can improve upon this foundation.

**Table 1.**
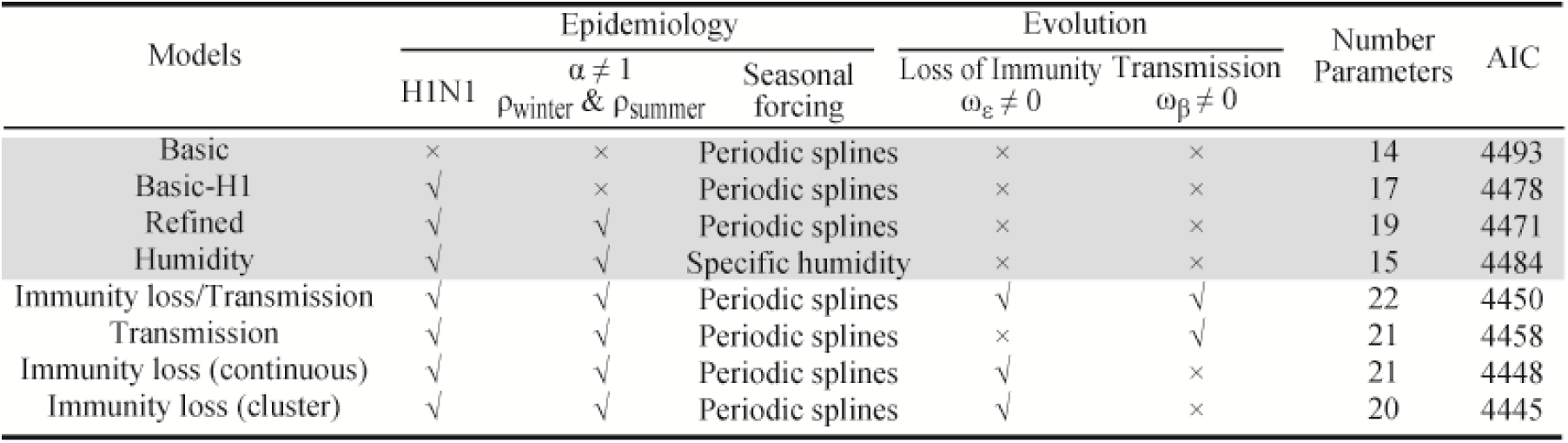
Model comparison. See Figure 1 for a diagram of the general epidemiological model. The ‘basic’ model is the well-known SIRS model with the transmission rate β (*t*) including only the seasonal forcing function and the loss of immunity rate ε (*t*) set to a constant. The ‘basic-H1’ model includes *R* _*H*1_ as an additional class to incorporate the loss of susceptible due to previous infection by H1N1, and use the observed incidence of H1N1 as a covariate determining the flux of individuals into this class. The ‘refined’ model additionally considers a different reporting error for the summer and winter seasons, and allows for the sub-exponential growth of the epidemic curve with an exponent α (smaller than 1) on *I* in the force of infection. The ‘humidity’ model replaces the spline-based seasonal forcing function with a humidity-based function. The next series of models includes the influence of the evolutionary index *E*(*t*) on the transmission rate β (*t*) and/or the loss of immunity ε (*t*). Among these evolutionary models (the ‘immunity loss/transmission’ model, the ‘transmission’ model, and the ‘immunity loss’ model), the best model (the ‘continuous’ model, compared to models without evolutionary compartment) based on the Akaike information criterion (AIC) is the one that incorporates a continuous dependence on *E*(*t*) only in the loss of immunity. A further improvement is achieved by the cluster model which incorporates evolutionary change as a discrete event for seasons with an antigenic transition (the ‘cluster’ model). AIC was calculated as: *AIC* = 2*k*-2*ln* (*L*), where *k* is the number of parameters and *L* is the maximum likelihood. The likelihood ratio test was used for model selection. Based on p values (smaller than 0.05), the basic-H1 and refined models are significantly better than the basic model, and the refined model is in turn significantly better than the basic-H1 model. Models with evolution, include the cluster model, are significant better than those without it (shaded). Among models with evolutionary information, the continuous and cluster model is significant better than the transmission model.

We measure antigenic innovation or evolutionary change of the virus at a given time (relative to a window of time in the past) using a novel evolutionary index readily computed from available sequences. The idea is to use amino-acid sequences to quantify change of the virus’ antigenic properties relative to those the human population has recently experienced. We therefore focus on sequences that encode epitopes known to be important for antibody binding and in which antigenic evolution is commonly observed (*14, 38, 39*). We define the index as a weighted average distance between the current virus to its predecessors in the past. So that more recent sequences are weighted more heavily than older ones, the weights are taken to be a decaying function of the inter-sequence interval. Figure 1C illustrates the estimated index. Since evolutionary novelty increases the probability that a virus escapes existing protective immunity and thereby achieves more efficient transmission, we incorporated this index as an external driver of the SIRS model, by allowing it to modulate either the duration of immunity or the transmission rate, or both. The model that includes evolutionary change in duration of immunity was best able to explain the data (the ‘continuous’ model) (Table 1). Simulations with the estimated parameters further demonstrate the improved performance of this formulation, with the median incidence reproducing the main trends in the interannual variation (Fig. 2B). As a general result (Table 1), the models that incorporate evolutionary information are significantly better than those that do not. Among the former, the models incorporating evolutionary change just in the immunity loss, or in both this parameter and transmission (the ‘immunity loss/transmission’ model) are comparable to each other, but perform significantly better than that with an effect only in transmission (the ‘transmission’ model). A degree of similarity between the two best models is to be expected in view of the fact that these two epidemiological parameters determine the overall infection rate so that it is difficult to disentangle an effect on the number of susceptibles from one on the per-susceptible risk of infection. At any rate, our results here indicate that inclusion of the evolutionary driver as a modulator of the duration of immunity is sufficient.

Our best model so far included a smoothly changing measure of evolutionary change based purely on virus genotype. It is recognized, however, that the genotype-phenotype map for antigenic properties of the virus is discontinuous, such that virus strains cluster antigenically and switches between clusters affect the population dynamics and phylodynamics of H3N2 in punctuated fashion (*4*, *10*). In particular, recorded antigenic cluster transitions are consistently followed by larger outbreaks (Fig. 1A). We found that our estimated index of evolutionary change followed observed antigenic cluster transitions: higher index values usually preceded a winter season with an antigenic cluster transition (compare Fig. 1A and Fig. 1C). This observation and what is known about the genotype-phenotype map of the virus (*4*, *40*, *41*) led us to a second evolutionary change index based on cluster transitions. In this ‘cluster’ model, the effect of a cluster transition is punctuated and localized in time: the rate of immunity loss is varied only during the summer season that precedes the winter season with an antigenic cluster transition. Specifically, the rate of immunity loss only during that time becomes a function of the degree of evolutionary change; with this change now measured by comparing current virus sequences to those two years ago, a time scale characteristic of cluster transitions (*4*). The resulting model performs better than the best model with continuous evolutionary change (Table 1 and Fig. 2C), although the most significant difference is between purely epidemiological models and those that incorporate evolutionary information (Table 1).

With our best model, the cluster model, we now turn to the task of predicting H3N2 incidence before the influenza season begins. For this purpose, we divide the data into a ‘training’ section (2002-2011) used to fit the models and an ‘out-of-fit’ one (2011-2016) used to evaluate their prediction accuracy. The implementation of the model for forecasting purposes requires particular assumptions on the drivers in the system since these observed quantities will by definition not yet be available over the time windows we wish to predict. For H1N1 incidence, we make the simplifying assumption that monthly averages for this quantity over the training set provide a sufficient approximation. Additionally, for the cluster model, an upcoming season dominated by a novel antigenic cluster needs to be anticipated in the summer before the transmission season. For this, we developed and tested a rule based on a published genotype-phenotype map for prediction of new antigenic variants. Specifically, when the proportion of antigenic variants accumulated during the summer season exceeded a given threshold, we took this to be predictive of a cluster transition in the following winter season (Fig. S2/S3; see Materials and Methods for details). Figure 3 shows the resulting retrospective predictions together with observations for each of the last five influenza seasons from 2011/2012 to 2015/2016. Two criteria were used to quantify prediction accuracy. The first compared the absolute monthly observed incidence to the medians of monthly predicted incidence from 1000 simulations. Predictions and observations are significantly correlated (*r* = 0.87 and *r* ^2^ = 0.76 for the monthly data; *r* = 0.95 and *r* ^2^ = 0.91 for the seasonal data; see Fig S4 for the data that include the most recent 2016/2017 season). Moreover, the observations mostly fell within the 97.5% confidence intervals (Fig. 3). Although the models tend to under-predict the absolute value of peak incidence, they do capture the overall interannual behavior of the trends reflected in both low and high seasons. The second criterion evaluates the model’s ability to predict an outbreak season by computing the probability of surpassing a selected incidence value deemed high by public health practitioners. We are interested here in evaluating the risk of an anomalous ‘high’ season relative to a typical season in the past and relative to a threshold level of cases of interest to public health. We consider first a threshold equal to the 50% quantile (median) of seasonal totals observed over the training dataset. A given flu season is forecasted as high or low risk level depending on the proportion of simulations that exceed the median, with the critical proportion that separates low and high levels chosen based on Receiver Operating Characteristic (ROC) curves (Fig. S5). Specifically, we predict a high risk level when more than 40% of the 1000 simulations surpass the median. All five seasons were predicted accurately based on this criterion (Table 2).

**Table 2.** H3N2 risk level forecasts for the US based on the cluster model. Seasonal risk level for H3N2 influenza virus is defined as high or low for each season from the out – of-fit period (2011-2017) compared to a reference level defined as the 50% quantile of the seasonal total H3N2 incidence cases in the corresponding training dataset. We defined an observed season as H3N2 high risk, when the observed total H3N2 incidence surpasses the reference level; and a H3N2 low risk season otherwise. For the forecasts, the percentage of 1000 simulations that exhibit a H3N2 high risk was obtained. When this percentage exceeded 40% (chosen based on Fig. S5), we forecasted a H3N2 high risk season. Otherwise, a H3N2 low risk season was predicted.

| Seasons | Observed | % High<br>(1000 simulations) | Forecasts<br>(>40%: high) |
| --- | --- | --- | --- |
| 2011/2012 | Low | 8.2 | Low |
| 2012/2013 | High | 99.6 | High |
| 2013/2014 | Low | 3.6 | Low |
| 2014/2015 | High | 99.9 | High |
| 2015/2016 | Low | 7.0 | Low |
| 2016/2017 | High* | 100.0 | High |

\* Based on the updated data from the weekly US influenza surveillance report until week 14 ending on April 8, 2017

**Figure 3.**
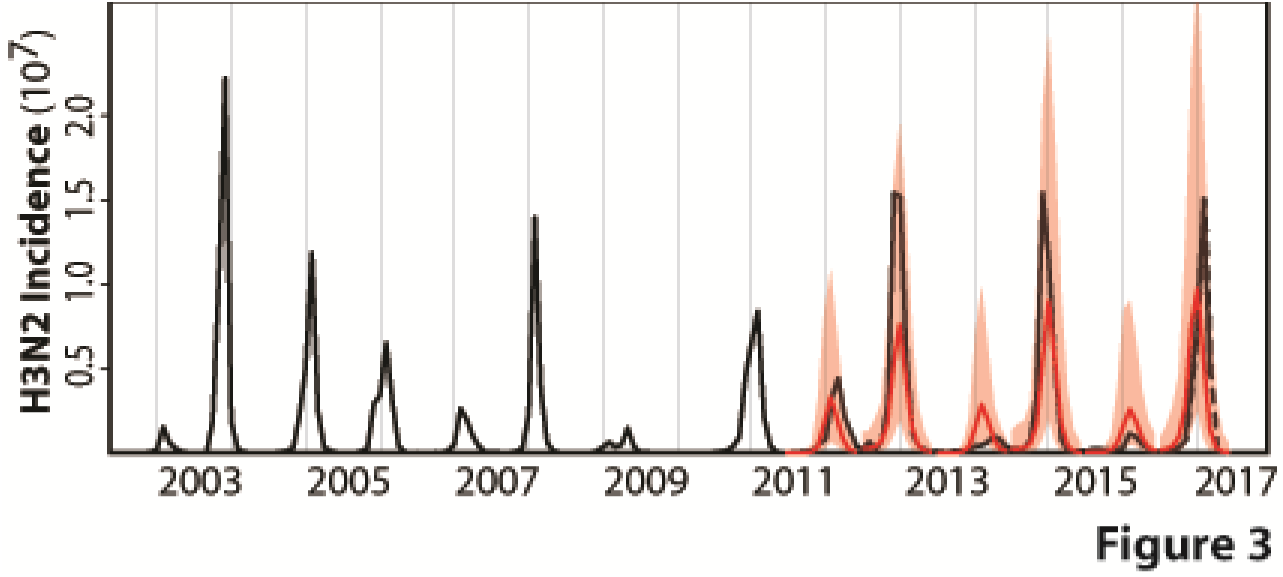
H3N2 incidence forecasts based on the cluster model for the US. Both retrospective forecasts (for each influenza season from 2011/2012 to 2015/2016) and a real forecast for the coming 2016/2017 influenza season are represented. These forecasts are simulated on a seasonal basis from estimated initial conditions starting in June and based on parameters estimated with all the data up to that point in time. The average monthly H1N1 incidence from this training dataset was used for forecasting purposes as the observation of this driver quantity would not be available. Similarly, the quantities specifying the evolutionary change of the virus was extrapolated as the sequences required for their computation would not be available. The black curve is the monthly observed H3N2 incidence; the red curve is the predicted monthly median incidence with shaded 2.5-97.5% quantiles from 1000 random simulations with the best models. The cluster model captures the occurrence of low and high seasons and forecasts high H3N2 incidence risk level for the 2016/2017 influenza season. The observed incidence data for the 2016/2017 influenza season, which were not yet available when this study was conducted, are shown with the dotted line (and based on data from the weekly US influenza surveillance report until week 14 ending on April 8, 2017).

To further evaluate prediction ability, we considered hindcast predictions, by removing one season at a time during the period from influenza season 2003/2004 to 2010/2011 and predicting its incidence, with the model parameters re-estimated each time based on the remaining data and exactly the same search strategy. Because there are multiannual correlations in the data, this test is less stringent and realistic than one based on multiple sequential out-of-fit seasons at the end of the time series. Nevertheless, it allows us to extend prediction evaluation and demonstrates high prediction accuracy (Fig. S6 and Table S1).

Encouraged by these results, we present a ‘real-time’ forecast prepared before the 2016/2017 influenza season and based on the data available up to June 2016, by the end of the 2015/2016 influenza season. Significant evolutionary change relative to previous viruses is indicated by our evolutionary index during the 2015-2016 influenza season (Fig. 1C), consistent with the observation that a number of antigenic variants were also accumulating over this period (Fig. S2). Concurrently, H1N1 dominated the 2015/2016 influenza season which would have resulted in an increased number of individuals susceptible to H3N2. The cluster model predicts that the risk level for seasonal H3N2 influenza should be high in the 2016/2017 influenza season for the US (Table 2), with a predicted annual incidence rate of 0.11 ([0.07, 0.15] for 2.5%-97.5% quantiles) (Table S2), consistent with the available observations (Fig. 3 and Table 2).

Forecast results based on the continuous model also capture the interannual trend in the size of epidemics (Fig. S7), and correctly predict risk levels above the 50% epidemic thresholds for the period between 2011 and 2017 (Table S3). The quality of the forecasts is lower however than that obtained with our best model (the cluster model) (compare Fig. 3 and Fig. S7). We also note that the 2016/2017 season is correctly predicted as high risk but that its peak size is over-estimated and its timing is earlier than that observed (Fig. S7). Finally, to further test the general approach, we applied it to a chosen region of the US-Department of Health & Human Services (HHS) region 3 (see Materials and Methods for details). A robust result is that the models with evolutionary information are significantly better at capturing the dynamics of seasonal H3N2 influenza than those without it (Table S4), although which particular evolutionary index is best can differ. Forecasts based on the best cluster and continuous models capture both the interannual variation of the outbreaks and disease risk for this US region (Fig. S8 and Table S5).

## DISCUSSION

Our results demonstrate the feasibility of improving epidemiological forecasts by incorporating information on evolutionary change into mechanistic models. Comparisons between models with and without this information show significant differences in their ability to capture the interannual variation in incidence data (Table 1), which underscores the importance of evolutionary change in the epidemiological dynamics of seasonal influenza. Our best models are able to capture the temporal behavior of observed incidence for H3N2 in the recent past in the US, and they provide the means to lengthen the lead time of prediction so that effective forecasts can be based in the summer, before the transmission season begins. Thus, this approach complements within-season forecasting efforts (*22*-*25*) and further informs public health preparedness. Earlier forecasts of incidence dynamics can aid public health efforts by indicating when to expect a surge in demand for healthcare resources and infrastructure. They can also contribute to the development of control strategies that take risk levels into consideration.

The use of evolutionary drivers in epidemiological dynamics follows an earlier study by of Axelsen and colleagues on long-term ILI incidence prediction in Tel Aviv, Israel (*42*). Their model incorporated the timing of known discrete antigenic changes in seasonal influenza, and demonstrated the importance of considering these discrete antigenic jumps and their interaction with the waxing and waning of immunity levels in the population. Prediction of multiannual temporal patterns over multiple seasons was shown possible after the observation of such an event and as long as another one did not recur, which is an impediment to real-time forecasting. A number of more mechanistic models coupling evolution and transmission dynamics have also been developed to address theoretical questions on the phylodynamics of seasonal influenza (*6, 7, 10, 12*). Because these individual-based, stochastic formulations are high-dimensional and computationally expensive to work with, they are not well-suited for the repeated estimation of parameters from time series data on reported cases or for epidemiological prediction. We have sought to construct much simpler epidemiological models suited for parameter inference based on surveillance, and for assimilating new data recurrently. Another approach to predict specific H3N2 incidence is based on its correlation with antigenic change as measured by the hemagglutination inhibition (HI) assay (*43*). Our approach exhibits a higher (Pearson) correlation between seasonal observations and predictions (0.83 for the whole US dataset from 2002 to 2016 and 0.90 for the testing dataset from 2011 to 2017, compared to 0.52 between antigenic change and H3N2 incidence for the period between 1998 and 2009) (Fig. S4B and Fig. S9).

Here, we have shown that readily available sequences of the virus can be used to construct an evolutionary covariate in both continuous and discrete versions. In the continuous model, the time scale of virus antigenic evolution is relatively short, with an average effective time of about 16 months (Fig. S10). At the same time, the estimated duration of homotypic immunity is relatively long: 30 years or even longer (Fig. S11). The importance of incorporating the short-term changes emphasizes the critical role of timely virological surveillance for identifying new emerging variants. Another intriguing observation related to the continuous measure of evolutionary change applied in our model is that the H1N1 pandemic of 2009 coincided with a valley in H3N2 fitness (Fig. 1C). This suggests that such times may provide a window of opportunity for the emergence of new types (including for cluster transitions of H3N2 itself). Thus, our evolutionary index, together with the proposed method for identifying and anticipating antigenic cluster transitions, could provide useful complements to the current surveillance system.

Our models made several simplifying assumptions, which can be investigated further and used to improve the approach in the future. The compartmental model did not include age structure (*44*) or social structure (*45*), proper inclusion of which might help to correct the underestimation of incidence peaks in this study. The evolutionary indices mainly consider antigenic change based on mutations in HA; these measures could be improved by further knowledge of antigenic phenotype (*46*, *47*), including other viral segments like protein neuraminidase (NA) (*48*). Also, other factors affecting the fitness of the virus could be considered (*14*, *49*), including receptor binding ability related to cell entry and transmission (*50*). Vaccination information could also be incorporated in the population dynamics. We decoupled the two-way interaction between subtypes H1N1 and H3N2 by including the effect of the former as a driver on the population dynamics of the latter. Although we observed that the dynamics of H3N2 was most strongly determined by its own evolutionary change, a more realistic model incorporating interactions between H3N2 and H1N1, and perhaps type B, could be considered (*51*). Our model can also be applied at finer spatial resolution and to other regions, especially in Asia, where the likely source of evolutionary novelty for the seasonal influenza virus is to be found (*5*, *52*). Preliminary investigation indicates that the general framework could be used in capturing and forecasting regional population dynamics of seasonal H3N2 influenza in US. Since immigration and emigration are also important processes in determining the local dynamics of seasonal influenza (*53*-*56*), a further step would consist of coupling regional dynamics to represent the effect of movement and the dependencies between adjacent regions. Similarly, at a larger scale one could incorporate information on global influenza circulatory patterns into the model. Finally, ways to better extrapolate the evolutionary covariate itself beyond the summer should be addressed. Overall, the fact that incorporation of pathogen evolution into epidemiological models increases forecasting skill should embolden future efforts to further improve on the model presented here.

The limits to lead times in influenza prediction are not set by chaotic dynamics as is the case for the weather system; they are determined instead by the stochastic nature of virus evolution. Formulating ways to take advantage in epidemiological prediction of the increased availability of genetic sequence data in surveillance efforts around the globe, is a promising area. One key limitation identified in our work is the low and variable number of sequences outside the transmission season, when this information would be most critical. Improvements to the general idea presented here will result from current efforts on purely evolutionary forecasting, which can provide better means to quantify antigenic change of the virus (*13, 14, 17, 40, 46, 47*), and to lengthen lead times further by concatenating evolutionary and evo-epidemiological prediction. Similarly, increased understanding of the virus’ genotype-phenotype map will also further inform this kind of effort. Ultimately, in the same way that routine weather forecasting provided the impetus for much better sampling of the climate system, incidence prediction is computationally feasible but will ultimately depend on the quality, depth and resolution of epidemiological and genetic surveillance.

## MATERIALS AND METHODS

### Data

HA Protein sequences of seasonal H3N2 influenza virus from US were downloaded from the Global Initiative on Sharing Avian Influenza Data (GISAID) (*57*). Sequences were then aligned with MUSCLE v3.7 using default settings (*58*). Undetermined amino acids were replaced by gaps, and only the HA1 domain was retained for further analysis. Outliers based on a reconstructed phylogenetic tree using FastTree 2 (*59*) with default settings, were manually removed.

Outpatient illness surveillance data and viral surveillance data were downloaded from FluView of the US Centers for Disease Control and Prevention (CDC) (*60*). Outpatient illness surveillance data includes information on patient visits to health care providers for ILI, which is collected through the US Outpatient Influenza-like Illness Surveillance Network (ILINet). The percent of patients presenting with ILI among all patient visits each week were used as indication of ILI in the US population. Viral surveillance data, including weekly influenza positive rate and subtype specific percentage data, were both from the US World Health Organization (WHO) Collaborating Laboratories and National Respiratory and Enteric Virus Surveillance System (NREVSS) laboratories. Seasonal and pandemic H1N1 influenza were combined together as seasonal H1N1 influenza, and the two lineages of seasonal B influenza were combined as seasonal B influenza. Un-typed influenza-positive specimens were assigned to either H3N2, H1N1 or B according to their proportions from typed specimens. The final weekly subtype specific incidence was calculated as the product of ILI positive rate, influenza positive rate, subtype specific proportion, and population size. Weekly incidence data was then aggregated to monthly data. Only incidence data from October 2002 to June 2016 was used in this study to focus on a period long enough to inform inference of model parameters, but to avoid earlier periods for which the sampling effort of genetic sequences was considerably lower and without surveillance data for the summer season. Monthly national level population estimates for the US were downloaded from the United States Census Bureau (*61*). Specific humidity data for the US were obtained from the National Land Data Assimilation System Phase 2 (NLDAS-2) products (*62*). These primary measurements are provided on a 0.125 degree grid. National data were averaged over all grids for the monthly data.

### Epidemiological model

We used a compartmental SIRS model to follow the flow of the population in susceptible, infected (and infectious) and recovered classes for seasonal H3N2 influenza. The model is given by the following equations:

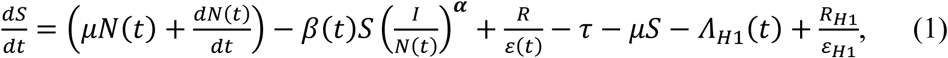

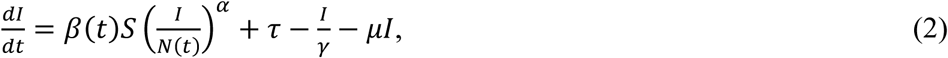

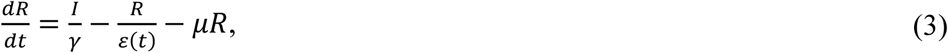

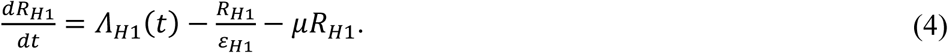

Where *S*, *I* and *R* denote the number of susceptible, infected and recovered individuals in the population, and *N*(*t*) is the population size at time *t*. The death rate *μ* was fixed to 0.015 per year (about 67 years lifespan). The total birth rate was quantified as 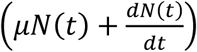 to reproduce the observed population increase over time. The exponent *α* is used to implement the nonlinear dependence of the force of infection on *I*, and the resulting sub-exponential growth of seasonal epidemics. τ is the external importation rate of H3N2 influenza cases which was fixed to 36.5 per year (1 import per 10 days) (*24*). The contact rate *β* (*t*) is given by:

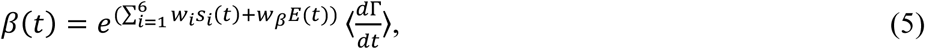

which includes three components: (*i*) seasonality implemented through six b-splines *s*_*i*_(*t*) with coefficients *w*_*i*_; (*ii*) evolutionary change *E*(*t*) (see below for details) with coefficient *w*_*β*_; (*iii*) and environmental noise simulated by a gamma distribution Γ (*63*). Under this model, the basic reproductive number is given by:

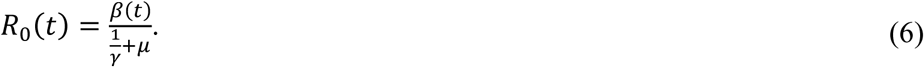

For the humidity-forced model, *β*(*t*) is given instead by the following expression based on (*21, 23, 24*):

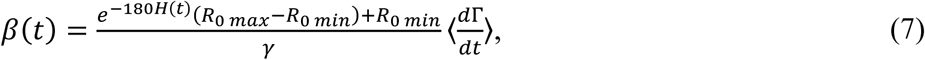

where *H*(*t*) is the specific humidity at time *t*, and *R* _0 *max*_ and *R* _0 *min*_ denote the maximum and minimum basic reproductive numbers and the basic reproductive number is here given by (*21, 23, 24*):

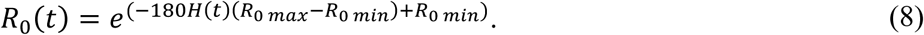

*ε* (*t*) is the average latent period at time *t*, given by:

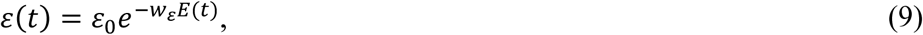

where *ε* _0_ is the basic latent period and *w* _*ε*_ is the scaling factor. *γ* is the average infectious period. An additional *R* _*H*1_ class was designed to track the reduction in susceptibles for H3N2 influenza due to cross-immunity, and therefore the protected population due to infection by seasonal H1N1 influenza. The rate of susceptible individuals temporarily moved to the *R* _*H*1_ class was measured by:

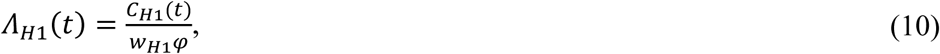

where *C* _*H*1_(*t*) is the observed incidence due to seasonal H1N1 influenza, *φ* is the reporting rate for H3N2, which is scaled here for H1N1 with the factor *w* _*H*1_. The average latent period for individuals in the *R* _*H*1_ class returning to the susceptible class is denoted by ε _*H*1_.

A measurement model is implemented that transforms the incidence in the transmission model to the actual observations by the reporting system. Specifically, reported cases were sampled from a normal distribution such that

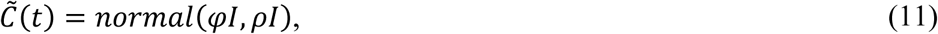

where *φ* denotes the reporting rate and defines the mean of observed cases, and the factor *ρ* defines the standard deviation as proportional to the size of the infected population. In addition, we impose the condition:

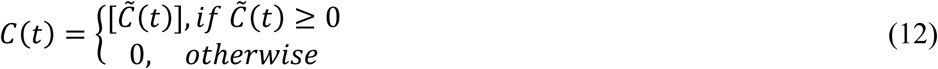

We note that the parameters of the measurement error model are fitted as part of the inference process. This is important since the value of the reporting rate can often be confounded with the degree of population immunity. As a result, we constrained its value to remain under 1, but also obtained a profile likelihood for this key parameter (Fig. S12).

### Evolutionary index

Two quantities were formulated to incorporate information on virus evolution into the epidemiological model. The first one, which varies continuously in time, is described here; the second, which varies discontinuously to reflect the punctuated antigenic change of the virus, is described below under ‘Antigenic cluster transitions and discrete evolutionary index’.

An evolutionary index, *E*(*t*), was used to measure the degree of evolutionary change of the virus at current time *t* (in months) compared to historical strains in the past. This index is therefore formulated as an weighted sum of normalized distance between current viruses and those in the past for amino-acid sequences encoding epitopes of the hemagglutinin protein on the surface of the virus (that is, parts of these protein recognized by the immune system in its antibody response). To begin, we can write the general expression:

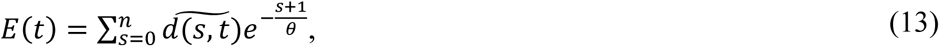

where 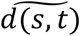 denotes a normalized distance between sequences at month *t* and those in previous season *s* back in time for epitope regions of HA1, and *n* is the total number of previous seasons including the current one for which we set *s* = 0. Thus, here, *n* = 19 and *t* = 1, … *l* (where *l* is the length of the incidence time series in months starting from October 2002). For the US, we defined the influenza season *s* as starting on July 1^st^ of one calendar year and ending on June 30 ^th^ of the following calendar year. Also, we refer to the summer season for the period between April 1^st^ and September 30 ^th^, and to the winter or ‘transmission’ season for that between October 1^st^ and March 31^st^. In the formula for *E*(*t*), changes relative to more recent viruses circulating in the population have a stronger weight than those relative to earlier viruses, and this weight decays exponentially back in time (scaled by θ in equation 13). Only a total of twenty years were considered to make sure there were no data missing for calculating *E*(*t*) of each month starting from October 2002. Distances 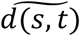 for a given month *t* were calculated relative to previous seasons (and not individual months) in the past because fewer viruses are typically sequenced during the summer season due to lower levels of incidence and associated weaker surveillance efforts. Also, early previous years exhibit multiple months without any reported sequence due to weaker sampling and sequencing efforts. Months and previous years were assigned based on date information that is at least monthly for sequences after 1992. For the earlier period between 1982 and 1992, although there are enough sequences for our calculations (described below), most of them lack detailed monthly information. As a result, approximate previous season assignments were made based on the reported calendar year (with calendar year 1990 assigned to season 1989/1990 and so on). Finally, in the formula for *E*(*t*) we sum these distances over time after weighting them back in time with an exponential decaying factor whose time scale is defined by the parameter *θ*.

The biology behind *E*(*t*) relates to the immune memory or protection existing at a given time in the human population for a new virus: the more similar this variant is to viruses in the past, the less likely it will be to infect people, since a higher probability exists that antibodies induced by viruses from previous infections will bind to it and stop the infection. Thus, the idea of a sum of weighted distance in sequence space is that of a quantity reflecting the movement of the virus away from variants the human population has been exposed to in the past. We include a decay function (controlled by a parameter *θ* that needs to be estimated) so that distances to more recent viruses have a higher weight when computing this average, given a time decay of antibody-mediated immunity. In other words, movement away from more recently observed antigenic variants would result in a higher evolutionary index and reflect a virus that is more novel from the perspective of the immunity landscape in the current human population.

We recognize that the weights can implicitly reflect additional processes that are not explicitly represented in the model, including the complex interaction of the age structure of the infected population (and contacts) and the effects on immune memory of age of exposure. The proposed quantity is intended to measure with a simple expression how much the virus has moved away from its recent predecessors in the sequence space that to the best of our knowledge reflects changes in the phenotype of interest. We note that since the rate of the decay backwards in time is one of the parameters inferred as part of fitting the overall model to the incidence time series, a possible outcome is for the decay to be negligible. In that sense, the inference process (and not an a priori assumption) determines the relevant time extent over which to evaluate the change in the virus. Although the idea is simple, the actual computation requires a series of steps because of geographic and temporal biases in numbers of sampled sequences, and the consideration of months and seasons in computing distances as we explain below.

First, an average distance *D*(*s*, *t*) was calculated based on 1000 distances (d_*m, st*_) among 1000 random pairs of sequences sampled from month *t* and previous season *s*. The actual value of d_*m, st*_ is calculated as the number of amino acid differences for epitope regions of HA1 (*38*). For distances to the current season (*s* = 0), distances were calculated based only on comparisons to sequences from earlier months. To avoid geographical and temporal sampling biases, we followed the practice of subsampling each random pair of sequences from sequences in both month *t* and previous season *s*, respectively, with equal probability from different states and from different months (or different previous seasons for earlier period before 1992) (*13*, *14*). This subsampling process was repeated 1000 times to get a mean value:

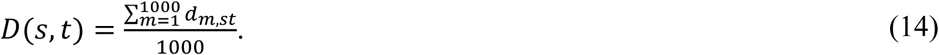

We note that *D*(*s*, *t*) is a matrix whose rows are previous seasons (from *s* = 0 in row 1, to *s* = 19 in the last row) and whose columns correspond to the months of interest in the time series of incidence to be analyzed (starting with October 2002 in column 1). We now proceed to normalize the entries of this matrix for each row, to correct for the effect of the passing of time within each season, which introduces an artificial trend in the unnormalized metric. Specifically, we normalize each term of the matrix by a mean value as follow:

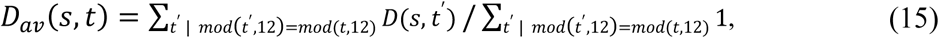

where the numerator sum is over all entries of the given row *s* that fall in the same month (as specified by the condition using the modulo operation *mod*), and the denominator sum simply counts the number of corresponding months. We then normalize the distances to obtain *d*(*s*, *t*):

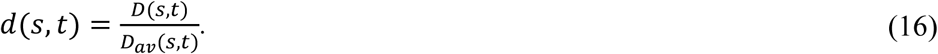

Finally, *d*(*s*, *t*) was interpolated (for months without sequences after October 2002, including March/May/July in 2004, May/July/September in 2005, May/June/July in 2006, December in 2009, February in 2010 and June in 2011) and smoothed by a cubic smoothing spline at a monthly scale (using the smooth.spline function in R package stats which uses a leave-one-out test to determine the smoothing parameter) to calculate 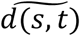 in equation 13.

### Antigenic cluster transitions and discrete evolutionary index

Antigenic cluster transitions were identified based on influenza season summary reports for seasons between 2003 and 2013 (*64*), and Morbidity and Mortality Weekly Reports (MMWR) for seasons between 2013 and 2016 (*65*-*68*), from US CDC. An influenza season summary report is produced after a season based on all available data for that season. In the report, CDC antigenically characterizes influenza viruses received in the past season, and assigns them to different groups based on antigenic similarity to the previous vaccine strain and also to a new vaccine strain. Here, different vaccine strains represent different antigenic clusters. We defined here an antigenic cluster transition as a season when more than half of the viruses were antigenically similar to the new vaccine strain. We note that the influenza ‘season’ used by US CDC is not necessarily the same as the one used in this study, but is instead variable often covering the transmission season. But because the vast majority of the data are from the transmission season, we can still apply an antigenic cluster transition to its corresponding influenza season, as defined in this study. Based on this criterion, antigenic cluster transition seasons are 2003/2004, 2004/2005, 2007/2008, 2009/2010, 2012/2013 and 2014/2015. For the 2006/2007 influenza season, although there was a change in vaccine strain, most of the circulating strains were not antigenically similar to the new vaccine strain (*64*). As a result, we considered that there was no antigenic cluster transition for the 2006/2007 influenza season. For the 2013/2014 influenza season, although there was a change in vaccine strain from A/Victoria/361/2011 to A/Texas/50/2012, the vaccine strains were antigenically similar (*68*). Thus, here too, we do not consider this vaccine strain change as an indication of an antigenic cluster transition.

For the model that incorporates evolutionary change through cluster transitions, we need to identify cluster transitions based on sequences, and when one such transition is identified, to quantify the degree of the change, which will influence model parameters (rate of immunity loss here) in the form of a step-function. That is, in the cluster model, an effect of evolutionary change in the virus is only applied to the epidemiological parameters when a cluster transition is identified (or predicted), and at this time, a measure of the magnitude of the change in the virus relative to the recent past is used as a covariate. In practice when implementing the model for actual prediction, we need to first predict that a new cluster will emerge and become establish to affect the transmission dynamics during the transmission season. This first step is implemented with a cluster transition rule based on a genotype-phenotype map for H3N2 previously published by one of us for both identifying and predicting antigenic cluster transitions purely based on sequence data without having to rely on antigenic assay data (*40*). (The latter concerns only a subsample of the virus circulating in the human population at a given time, and is typically available with a delay relative to sequence data). Although the genotype-phenotype map relies on sequences, it uses a number of properties of the hemagluttinin protein derived from the sequences including biophysical properties, and not just amino-acid distances at epitope sites. A brief description of this map and how it is used here for prediction purposes is included below in the section on Forecasts. Here, we note that this first step allows us to implement prediction of an antigenic cluster transition ahead of the transmission season, during the summer. Having identified (for retrospective data) or predicted (for ‘out-of-fit’ data/forecasting purposes) the emergence of a new antigenic cluster, we proceed in a second step to quantify how much the new viruses differ from those in the recent past.

We specifically define the following quantity to measure evolutionary change (at the monthly scale) for the month of September:

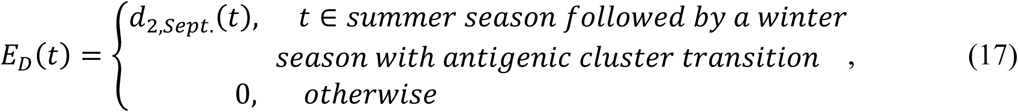

where *d*_*2, Sept*_. was calculated as a normalized distance based on epitope regions of HA1 between sequences for September of the current season and sequences of the previous second season (for example, *d*_*2, Sept*_ in 2000 was calculated based on sequences from September of current 2000/2001 influenza season and 1998/1999 influenza season, that is *s* = 2 in the notation introduced above). We chose September because a new antigenic cluster would need to be established before the winter season to effectively influence the dynamics of that transmission season. We chose a comparison between September and the previous second season (our ‘reference’ season) for two reasons: first, the average effective period back in time estimated with parameter θ for the evolutionary index is about two years (Fig. S10), and second, this period is close to the average time of cluster replacements (*4*).

Because there are typically a limited number of sequences for the summer season, distances calculated based on those sequences can vary considerably depending on sampling effort. Therefore, to obtain a final value of *d*_*2, Sept*_., we used a similar procedure than that used for calculating 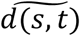 above. First, with the same subsampling procedure to address geographical and temporal sampling biases, we calculated an average hamming distance for strains in each month (from October of the previous year to September of the current year) and those from the corresponding reference season (*s* = 2). Then we normalized these values to calculate *d*(*s*, *t*), here *d*(*s* = 2, *t*) Finally, in order to lower the impact of stochasticity in the data, especially for data from summer seasons, *d*_*2, Sept*_. was computed as the fitted value (or extrapolated if missing) for September, based on a linear regression of distances *d*(*s* = 2, *t*) for months between October of the previous year and September of the current year.

For the purpose of fitting the time series data, the resulting *E* _*D*_(*t*) was introduced in the model as described in equation 9. When the goal is instead that of specifically forecasting ahead of the transmission season, we need to anticipate cluster transitions and the value of *E* _*D*_(*t*) accordingly. We describe the approach we take for this purpose in the section below on Forecasts.

### Parameter estimation

The resulting SIRS model was fitted to the data using Likelihood Maximization by Iterated particle Filtering (MIF) in the R package pomp (*69*, *70*). Both parameters and initial conditions (for *S*, *I*, *R* and *R* _*H*1_) were estimated based on the likelihood function:

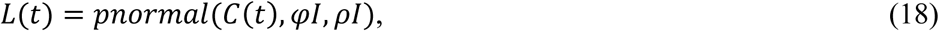

with *C* (*t*) > 0. If *C* (*t*) = 0, *L* (*t*) was set to a very small value equal to-10000 (log scale) as a penalty. The search of parameters and initial conditions was started with a grid of 10,000 random combinations sampled using the Latin Hypercube sampling (*71*) from wide ranges. This step was followed with additional phases of increasingly localized searches. Confidence intervals were estimated separately for each parameter with the target parameter fixed at different values while allowing estimation of all other parameters also using MIF (*69*, *70*). The Akaike information criterion (AIC) was used to measure goodness of a model (*72*). The AIC score takes into account model complexity and penalizes the likelihood based on the number of parameters. The likelihood ratio test was used for model selection for nested models (*73*).

### US regional models

For a test of robustness of our general framework, we applied it to data from one US region, the US Department of Health & Human Services (HHS) region 3, which includes Delaware, District of Columbia, Maryland, Pennsylvania, Virginia and West Virginia. The epidemiological, virological and population data were downloaded from the same websites as for the national data. We note that the regional data (ILI positive rate, influenza positive rate and type/subtype specific proportion) is sparser with frequent data missing during the low season. Because the regional data are therefore noisier, we need to smooth it for further use. We did so by smoothing the incidence data from peak (time point with highest incidence during a winter season) to peak by local linear regression using a 4-weeks window. We also slightly revised our model to make its fitting less sensible to the data during the low seasons, which are mostly interpolated. We did this by adding a constant (200) to the reporting error through ρ in equations 11 and 18 so that likelihoods calculated based on the low seasons vary less and contribute less in differentiating model performance. We also increased the importation rate from 0.1/day to 10/day to allows for the more frequent movement of people between regions within US (*53*, *55*). Additionally, we used the national sequence data for the evolutionary covariate, under the assumption that from the perspective of evolutionary change the whole country would be largely synchronized (*5*, *53*). The choice of spatial scale could be examined further in the future, although more limited sequences are available for the regional level.

### Forecasts

Forecasts of incidence dynamics were obtained for a given season through three steps. First, the cluster model was trained based on the dataset before the target season using exactly the same procedure described above. Second, the model with the best likelihood was chosen and used to estimate the initial conditions for the forecasts. These initial conditions require estimates of the ‘hidden’ (un-measured) variables in the model, *S*, *I*, *R* and *R* _*H*1_ in June of the year for which the predictions will be made. MIF allows one to estimate these variables as the filtered states of the system at that time given the observations of the cases. Third, forecasts were obtained through forward simulations for the target season with the selected model and starting from these initial conditions.

Because the system of equations in our models is non-autonomous, including external variables or drivers whose values must be specified independently from the dynamics, real forecasts (vs retrospective ones) require specifying assumptions for these drivers whose observation in the future is by definition unavailable. Specifically, we require information on both *C* _*H*1_(*t*) in Λ _*H*1_(*t*) and *d*_*2, Sept*_. in *E* _*D*_(*t*) for the simulations. First, for *C* _*H*1_(*t*), the average monthly value from the training dataset was used. This is a simplification and a rough approximation but a reasonable choice in the absence of modelling the coupled dynamics of H1N1 and H3N2. This approximation is most likely to be sufficient, when predictability of our models is evaluated, if the population dynamics of H3N2 is most strongly driven by its own evolutionary change rather than by the precise levels of H1N1 (our results suggest that this is indeed the case). Second, the evolutionary change between new and old clusters is also needed, and this quantity can be extrapolated linearly as the value for September using the same procedure for calculating *d*_*2, Sept*_. above, but based on sequences only from October of the previous year to June of the current year.

In addition and importantly, the weekly reports used to identify antigenic cluster transitions will not yet be available at the time of forecasting. Previously, a naive Bayes model was developed as a genotype-phenotype to translate sequence changes in HA to antigenic changes, and specifically calculate an odd ratio measuring antigenic similarity between a pair of strains given their sequences (*40*). Instead of only relying on the number of amino acid changes in epitopes of HA, this method employs several additional features that are related to intrinsically physiochemical mechanisms of antigenic change to predict the antigenic stasis of a strain variant (*40*). Based on this method, quarterly measures of the proportion of antigenic variants (PAV) were calculated here. This quantity provides the proportion of pairs that are antigenically different (odd ratio < 1) among all pairs between sequences of a specific quarter and those in the previous year. Again, the same subsampling process was applied, but based on time points for quarters not months.

To predict a cluster transition, we examined rules that combine a local increase in PAV with this quantity exceeding a threshold. This cutoff value was selected based on Receiver ROC curves (lower bound of best accuracy, see Fig. S3). Again, because a new antigenic cluster would need to be established before the winter season, we chose to evaluate PAV in the third quarter (July 1^st^ to September 30 ^th^) right before the coming winter season: PAV for the third quarter was linearly extrapolated based on data in the previous three quarters (first and second quarter of the current year and the fourth quarter in the previous year; that is from October 1^st^ to June 30 ^th^. The rule we constructed on the basis of PAV was meant to combine a requirement that there is sufficient novel viruses accumulating, at a time in which novelty is rising. Specifically, based on data from the current season, if PAV increases from the first quarter to the second quarter and crosses the selected cutoff of 0.11 for the third quarter (whose value was extrapolated as explained above), an antigenic cluster transition was identified for the upcoming season. If PAV decreases from the first quarter to the second quarter, but is still higher than the cutoff value of 0.11 for the third quarter (again, extrapolated), a cluster transition will also be assigned for the upcoming season, with the additional requirement that an antigenic cluster transition was not already assigned for the current season. Otherwise, no cluster transition will be anticipated for the upcoming season. With this rule and for data between 2002 and 2012, we would have correctly anticipated all antigenic cluster transitions (true positive rate equal to 1 and false positive rate equal to 0) with a cutoff value of PAV ranging from 0.11 to 0.2. The lower bound of 0.11 was chosen as the cutoff in this study.

For the continuous model, we also need to know *C*_*H*1_(*t*) and *E*(*t*). For *C* _*H*1_(*t*), the average monthly values of H1N1 incidence were used. *E*(*t*) was linearly extrapolated up to September using the data available until June of the current season, then kept constant.

When predicting the risk level (high or low) of a target season, we first define an epidemic relative to a reference threshold, defined initially as the median (50% quantile) of the seasonal total incidence in the training dataset. We then calculated the percentage of simulations above this threshold among 1000 simulations for the given target season. This percentage provides a probability of exceeding the given epidemic level. We can again use ROC curves and the training data set to establish which probability should be exceeded to predict a high risk. If the percentage is above a cutoff (upper bound of best accuracy), the target season is predicted with high risk; otherwise, it is a low risk season (see Fig. S5 for the US national data based on the cluster model, Fig. S13 for the US national data based on the continuous model, and Fig. S14 for the HHS region 3 data). Although a natural choice might be 50% of the simulations, ROC curves can indicate that lower percentages should indicate risk given the tendency of the model to under-predict the size of the peaks.

### Cross-validation (hindcast)

In order to test predicting ability further, and specifically for different patterns of alternating dominant subtypes in adjacent seasons, we conducted a cross-validation analysis for the training dataset itself covering the period from 2002 to 2011. For each influenza season from from 2003/2004 to 2010/2011, one at a time, the cluster model was fitted de novo by removing the target year and using the same search strategy than for the full data set before. With the resulting specific parameters, a prediction was generated using the same strategy described above and calculating mean H1N1 incidence with all years except the target one. In practice, the fitting of the model is implemented in MIF with parameters prevented from performing a random walk during the window of time that contains the target year (so that the corresponding data is not used in the filtering process), and with the likelihood evaluated by setting *L*(*t*) = 0 (in the log scale) in equation 18.

## SUPPLEMENTARY MATERIALS

Table S1-S5

Figs. S1-S14

## ACKNOWLEDGEMENTS

The authors acknowledge the Research Computing Center at the University of Chicago for providing computational resources for this research. AAK has been supported by the National Institutes of Health (1R01AI101155) and by MIDAS, National Institute of General Medical Sciences U54-GM111274. MP, AAK and XD conceived and designed the experiments. XD performed the experiments and analysis. All authors interpreted the results and wrote the manuscript. The authors have declared that no competing interests exist.

